# Regulatory Features and Functional Specialization of Human Endogenous Retroviral LTRs: A Genome-Wide Annotation and Analysis via HERVarium

**DOI:** 10.64898/2026.02.17.706328

**Authors:** Tomàs Montserrat-Ayuso, Aurora Pujol, Anna Esteve-Codina

**Affiliations:** Centre Nacional d’Anàlisi Genòmica (CNAG), Baldiri Reixac 4, 08028 Barcelona, Spain; Universitat de Barcelona (UB), Barcelona, Spain; Neurometabolic Diseases Laboratory, Bellvitge Biomedical Research Institute (IDIBELL), Barcelona, Spain; Centre for Biomedical Research on Rare Diseases (CIBERER), Instituto de Salud Carlos III, Madrid, Spain; Catalan Institution of Research and Advanced Studies (ICREA), Barcelona, Spain

## Abstract

Human endogenous retroviruses (HERVs), remnants of ancient retroviral infections, account for over 8% of the human genome and remain an underexplored source of cis-regulatory elements and protein-coding remnants. Here we present HERVarium, an integrated and interactive database that provides access to systematic annotations of the protein-domain architecture of internal HERV regions and the regulatory landscape of their LTRs across the human genome. Building on our previous catalog of conserved retroviral domains, we classified over 400,000 LTRs by structure (solo, 5′, or 3′ LTR), proximity to transcription start sites (TSS), potential transcription-factor binding-motif (TFBM) burden, and computationally reconstructed their canonical U3–R–U5 substructure, revealing that two-thirds retain recognizable segmentation, particularly those adjacent to conserved internal domains. LTRs flanking internal regions with conserved Gag, Pol, or Env domains tend to be longer and richer in motifs, suggesting coordinated maintenance of coding and regulatory potential. Conversely, solo LTRs positioned at gene TSSs exhibit promoter-like architectures enriched in motifs for developmental and proliferative regulators, while selectively lacking motifs associated with neuronal differentiation. Some 5′ LTRs at lncRNA TSSs also flank conserved retroviral domains, indicating that certain lncRNAs may derive from transcriptionally active, structurally intact HERV loci. HERVarium provides a comprehensive, interactive resource to explore and download these annotations at single-locus resolution.

## Introduction

Endogenous retroviruses (ERVs) are remnants of ancient germline infections by exogenous retroviruses that became permanently integrated into host genomes and are now vertically transmitted (Stoye 2001). In humans, endogenous retroviruses (HERVs) account for ∼8% of the genome(Griffiths 2001). Although most HERVs have lost infectivity through the accumulation of mutations, recombination, and deletions, they persist as molecular “fossils” that record the deep evolutionary history of retroviruses and their hosts (Bannert and Kurth 2006). These molecular fossils capture both the diversity of now-extinct viruses and the evolutionary arms race between retroviruses and vertebrate immune systems (Bannert and Kurth 2006; Srinivasachar Badarinarayan and Sauter 2021). Typically, HERVs are represented in the genome as degraded proviruses, solo long terminal repeats (LTRs), or fragmented internal coding regions (Bannert and Kurth 2006; Ueda et al. 2020).

LTRs are critical regulatory modules of retroviruses, originally providing promoters, enhancers, and polyadenylation sites required for viral gene expression (Griffiths 2001; Thompson et al. 2016). Even after the decay of coding capacity, many HERV-derived LTRs retain transcriptional regulatory activity and have been co-opted as cis-regulatory elements in the host genome (Thompson et al. 2016; Ito et al. 2017). Compared to other transposable elements (TEs), LTRs are particularly well suited for exaptation because they already encode complete promoter/enhancer modules (Thompson et al. 2016). Consequently, they have had a substantial impact on shaping transcriptional networks during primate evolution (Thompson et al. 2016; Jacques et al. 2013). Comparative genomics and epigenomic profiling indicate that transposable elements—including ERVs—have profoundly reshaped the primate regulatory landscape: up to 63% of primate-specific open chromatin regions derive from TEs, with ERVs being disproportionately represented (Jacques et al. 2013). Specific LTR families, such as HERVH LTR7, act as promoters in pluripotent stem cells (Zhang et al. 2022), while others contribute enhancers in immune and tissue-specific contexts (Thompson et al. 2016). Thus, LTRs contribute a rich but heterogeneous layer of regulatory diversity in the human genome.

Beyond cis-regulatory contributions, HERVs have been repeatedly co-opted at both the gene and regulatory level. At the coding level, the most celebrated examples are *syncytins*—retroviral envelope proteins recruited for placental development across multiple mammalian lineages (Cornelis et al. 2015). At the regulatory level, many solo LTRs have been adopted as promoters and enhancers of host genes, rewiring transcriptional networks in early embryogenesis or immunity (Thompson et al. 2016; Chuong et al. 2016). Large-scale ChIP-seq and motif analyses confirm that LTRs often harbor dense clusters of transcription factor (TF) binding motifs, including pluripotency factors (SOX2, NANOG, POU5F1), hematopoietic regulators, and immune-related TFs (Ito et al. 2017). Notably, co-opted LTRs are often enriched in motifs associated with pluripotency and early development, consistent with selective pressures favoring regulatory plasticity in progenitor and stress-responsive contexts (Thompson et al. 2016; Ito et al. 2017; Chuong et al. 2016).

The regulatory and coding potential of HERVs is double-edged. On one hand, repurposed elements contribute to normal development, immunity, and gene regulatory diversity. On the other hand, aberrant HERV activation has been linked to neurological disease (Küry et al. 2018; Duarte et al. 2025), cancer (Li et al. 2010; Zhao et al. 2011; Rycaj et al. 2015), and autoimmunity (Viret and Bynoe 2024). For example, HERV-W and HERV-K envelope proteins have been implicated in multiple sclerosis (MS) and amyotrophic lateral sclerosis (ALS) disease onset or progression (Küry et al. 2018; Duarte et al. 2025), while reactivated HERVs can facilitate protein aggregate spreading in neurodegeneration (Liu et al. 2023). These dual roles underscore the importance of distinguishing nonfunctional, background HERV activity from loci that preserve functional regulatory or coding potential.

Despite these potential impacts in human disease states, current resources for analysis and interpretation remain fragmented. Existing databases typically specialize in either broad structural catalogs of HERV loci (e.g., HERVd, which integrates internal regions and LTRs but lacks fine-grained domain or TF-binding site annotation (Pačes et al. 2002)), or in coding remnants (e.g., gEVE (Nakagawa and Takahashi 2016)), or in regulatory features (e.g., dbHERV-REs (Ito et al. 2017)), without integrating both perspectives. Moreover, classification of LTRs into structural categories—solo, 5′, and 3′—remains incomplete, limiting our ability to assess how regulatory activity relates to proviral structure, and to distinguish exapted LTR promoters from passive innocuous LTR fragments left behind by provirus decay.

In our recent work (Montserrat-Ayuso et al. 2026), we systematically annotated conserved protein domains across >120,000 HERV-derived ORFs in the human genome, uncovering a surprising abundance of intact retroviral-like domains—including nearly full-length *gag*, *pol*, and *env* sequences—in both young and ancient HERV families. These findings suggested that structural conservation can persist more widely than previously appreciated, but also highlighted a major gap: the regulatory landscapes of the flanking LTRs were not considered.

Here, we address this unmet need by systematically identifying potential transcription factor binding motifs (TFBM) across all HERV LTR sequences in the human genome and classifying each LTR as 5′, 3′, or solo, alongside annotation of internal regions as having only a 5′, only a 3′, both, or no LTRs. In addition, we computationally reconstructed the canonical U3–R–U5 architecture of each LTR, which reflects the classical organization of retroviral long terminal repeats: the U3 region contains the promoter and enhancer elements that control transcription initiation, R corresponds to the transcribed sequence repeated at both ends of the viral RNA, and U5 carries signals involved in reverse transcription and polyadenylation (Coffin et al. 1997). This integrative framework connects the structural context of HERVs with the regulatory architecture encoded in their LTRs.

To make these annotations accessible, we developed HERVarium, an interactive database and web application that integrates a built-in genome browser. The browser displays multiple HERV-centric annotation tracks—including internal regions, conserved protein domains, LTR elements, U3–R–U5 segments, PBS/PPT sites, promoter and PAS signals, and locus-resolved transcription factor binding motifs—together with GENCODE genes and optional ENCODE DNase-seq and GTEx RNA-seq coverage tracks. HERVarium enables locus-level exploration, cross-filtering, and visualization of regulatory and protein-coding features across the HERV landscape. The platform is available for local installation via GitHub (https://github.com/funcgen/HERVarium) and Zenodo (https://doi.org/10.5281/zenodo.18551738), and a public web server will be deployed by the time of publication.

Using this unified framework, we found that LTRs flanking internal regions with conserved domains are significantly longer and harbor denser clusters of regulatory motifs than solo LTRs, suggesting coordinated retention of structural and regulatory potential. Moreover, elements that retained a well-defined U3–R–U5 substructure also showed the highest correspondence with conserved internal domains, indicating that preservation of internal coding potential tends to be accompanied by greater maintenance of promoter and polyadenylation architecture. Solo LTRs overlapping gene promoters exhibited selective enrichment for early embryonic development-associated motifs, while being depleted of motifs linked to neuronal lineage commitment and differentiation. Together, these findings illustrate how HERVarium serves both as a comprehensive database and as a discovery tool, facilitating the exploration of the co-occurrence of structural conservation and regulatory specialization in HERV loci.

All underlying code and pipelines used for LTR classification, motif annotation, and database construction are openly available via GitHub (https://github.com/funcgen/herv-regulatory-map), ensuring reproducibility. In addition, a permanent Zenodo repository provides versioned access to the full set of LTR annotations (https://doi.org/10.5281/zenodo.17602210), paralleling the data release strategy of our previous HERV domain annotation resource. These open resources ensure that HERVarium can be readily integrated into computational and experimental workflows and continuously extended by the community.

## Results

### HERVarium resource overview and annotation framework

To enable a systematic investigation of the regulatory potential of HERV LTRs, we developed HERVarium, an integrated reference that unifies structural annotation of internal HERV regions with genome-wide regulatory annotation of their associated LTRs. Building on our recent map of conserved retroviral protein domains (Montserrat-Ayuso et al. 2026), we extended this framework to include all LTRs annotated in the human genome (hg38) and linked each element to nearby internal regions of the same subfamily and strand.

Each LTR was classified according to its genomic relationship with internal HERV regions, distinguishing solo LTRs, 5′ LTRs, 3′ LTRs, and ambiguous cases. This stratification separates LTRs likely to function as independently co-opted regulatory elements (for example, solo LTRs located at host-gene transcription start sites) from LTRs that remain embedded within proviral structures retaining internal coding potential. Integration with GENCODE v48 further enabled classification of LTRs by their distance to host-gene transcription start sites, providing a common coordinate system for downstream analyses of motif content, positional architecture, and gene associations.

Across all annotated LTRs, we quantified TFBMs using FIMO and integrated these data with the conservation status of linked internal regions. This joint annotation links regulatory motif content to proviral structural context, enabling comparative analyses of how motif burden and density vary with LTR architecture and internal domain preservation. In parallel, we reconstructed the canonical U3–R–U5 organization of each LTR and identified PBS and PPT in 5′ and 3′ LTRs, respectively. Together, these annotations capture the degree of retention of promoter- and polyadenylation-associated features across HERV families and structural contexts.

To facilitate exploration of this multi-layered annotation, HERVarium provides an interactive interface that allows users to filter and inspect individual HERV loci, visualize LTR structure and motif content, and examine loci in genomic context alongside internal domains and selected external datasets, including ENCODE DNase I hypersensitivity profiles and GTEx tissue expression tracks (Figure 1). This unified environment links proviral structure, internal domain conservation, and regulatory architecture at single-locus resolution, supporting both hypothesis-driven and exploratory analyses across the full repertoire of human HERV elements.

**Figure 1.**
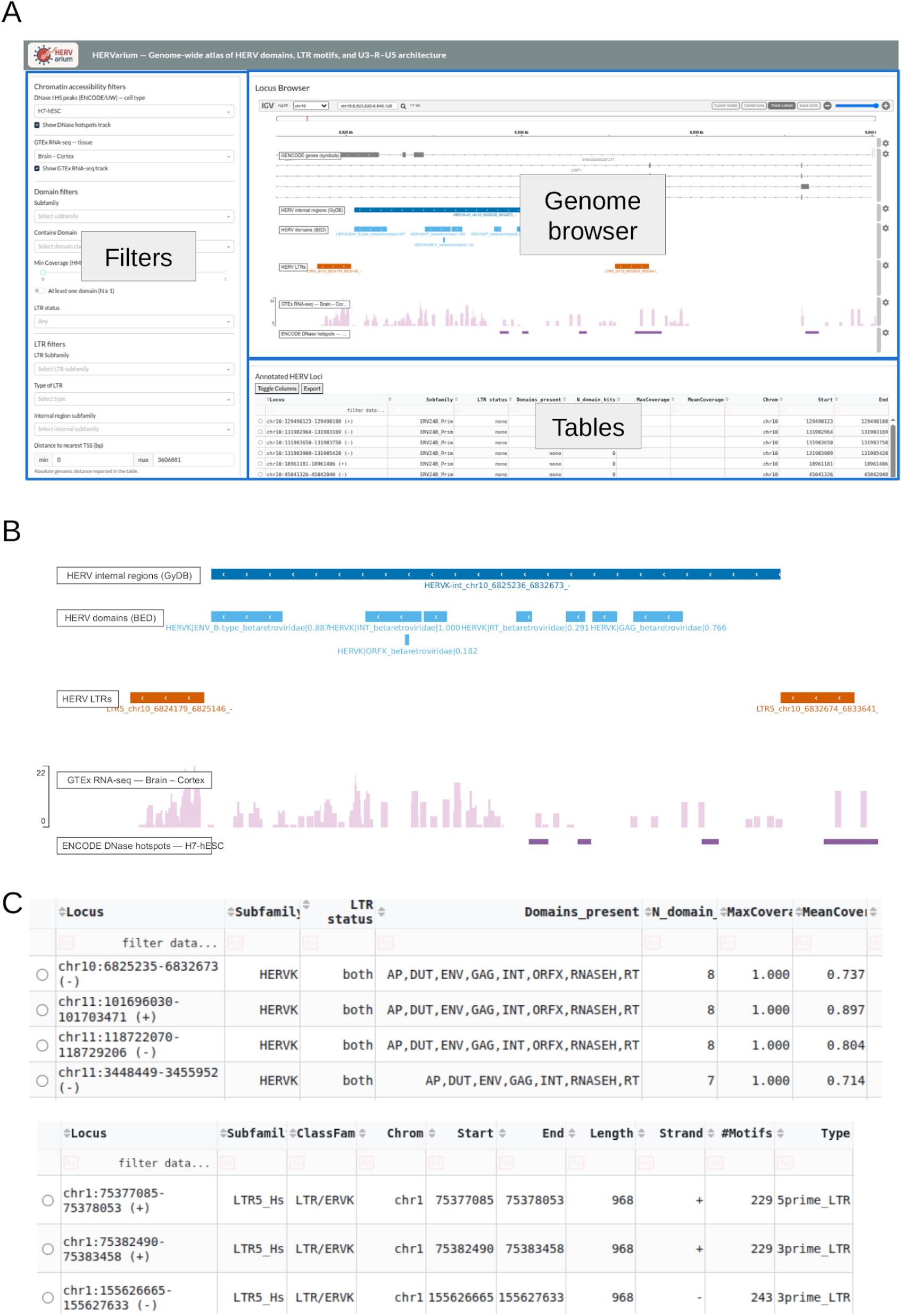
A) Screenshot of the HERVarium web interface illustrating the three main components of the application: interactive filter controls (left), an embedded genome browser displaying integrated HERV annotation tracks (center), and dynamically linked annotation tables (bottom). Filters allow selection by HERV subfamily, LTR structural class, internal domain conservation, motif burden, and U3–R–U5 features, with synchronized updates across all views. B) Genome browser view of a representative HERVK locus, showing integration of multiple annotation layers at single-locus resolution. Tracks include internal HERV regions, conserved retroviral protein domains, flanking 5′ and 3′ LTRs, transcription factor binding motifs, reconstructed U3–R–U5 segments, and associated PBS/PPT features (not all tracks shown), displayed together with optional external data tracks such as ENCODE DNase I hypersensitivity and GTEx RNA-seq coverage. C) Examples of HERVarium annotation tables reporting internal HERV loci (top) and linked LTRs (bottom). Tables summarize locus coordinates, subfamily assignment, LTR structural status, domain composition and coverage, motif counts, among other characteristics, and are directly linked to the genome browser to enable rapid navigation and comparative analysis across loci.

This annotation framework provides the foundation for the analyses below, which examine how LTR regulatory features vary with structural context, internal domain conservation, and genomic positioning.

### Conserved HERV LTRs Are Longer and Enriched for Regulatory Motifs

To explore how regulatory potential relates to structural conservation in HERVs, we mapped TFBM occurrences across all LTR sequences using FIMO and summarized them per LTR as motif burden (total matches) and motif density (matches per bp). LTRs were stratified by structural context—solo; 5′; 3′; or tandem (Figure 2A), the latter identifying LTRs that form part of a complete LTR–internal–LTR tandem. For LTRs linked to internal regions, we further separated loci based on whether the adjacent internal region exhibited high HMM profile coverage (≥0.95 for at least one protein domain; “Full”) or degraded/absent domains (“Deg”). LTR lengths were compared across the same categories.

**Figure 2.**
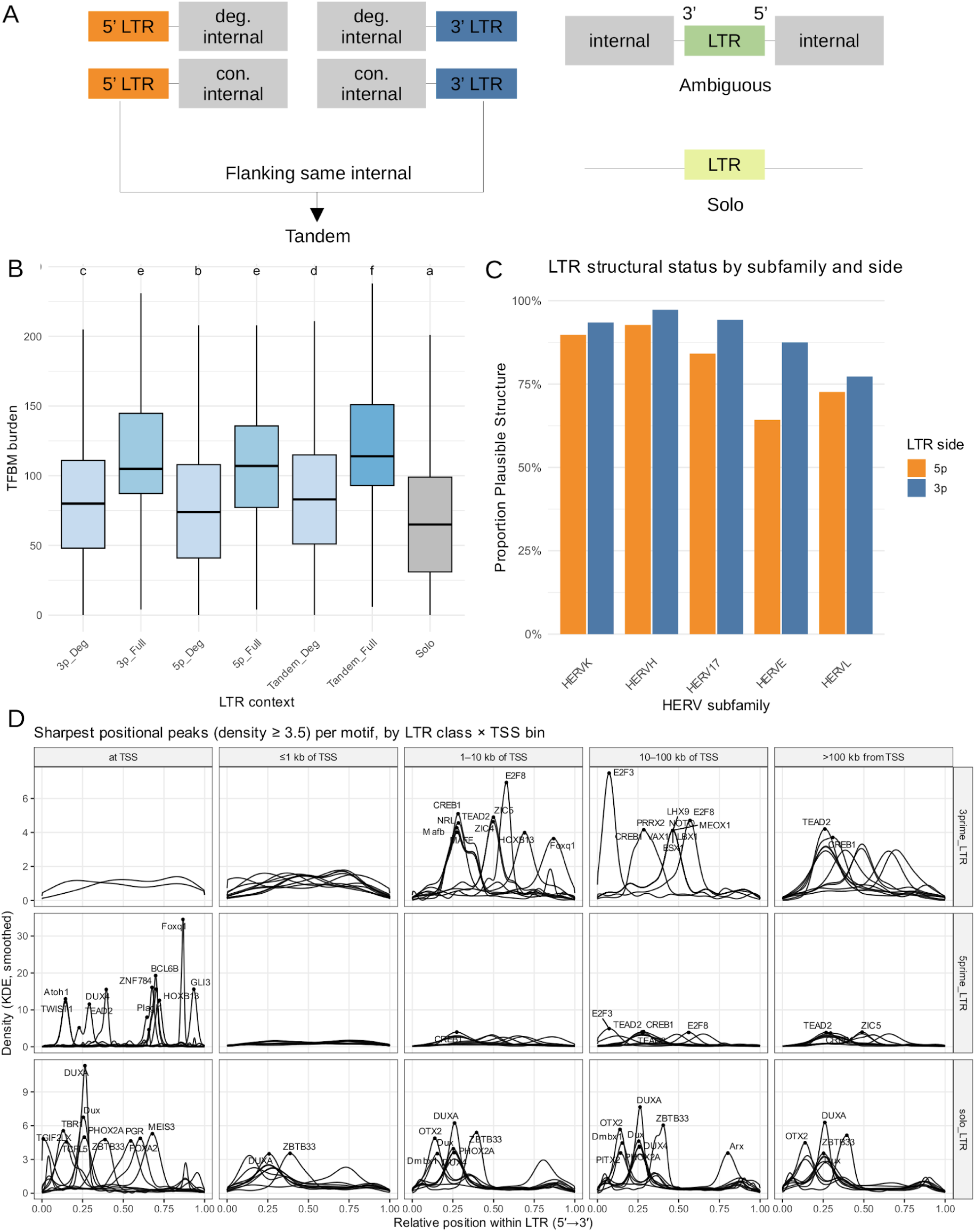
A) Classification of LTRs based on the conservation status of their linked internal regions. LTRs flanking the same internal region on opposite sides were assigned to the “tandem” category. Ambiguous cases—where a single LTR was linked to multiple internal regions with conflicting labels—were resolved by prioritizing links to internal regions with conserved (full) domains or to tandem configurations; remaining ambiguous LTRs were discarded from the TFBM burden analysis. B) LTRs flanking internal regions that retain at least one conserved retroviral domain displayed significantly higher TFBM burden than those flanking fully degenerated internal regions; solo LTRs contained the fewest TFBMs overall. Different letters above boxplots indicate groups that differ significantly (adjusted p-value < 0.05). C) Across HERV families, 5′ LTRs exhibited fewer well-defined U3/R/U5 segments, reflecting greater structural decay relative to 3′ LTRs. D) Several TFBMs exhibited strong positional enrichment at specific relative locations within the LTR. This positional signal was particularly pronounced in 5′ LTRs that overlap host-gene transcription start sites (TSSs). Moreover, the positional patterns differed markedly depending on genomic context, with LTRs located at host-gene TSSs showing distinct motif distributions compared with LTRs positioned far from TSSs.

Global differences were highly significant for all measures (Kruskal–Wallis: motif burden *p* ≈ 0; motif density *p* = 5.03×10^-108^; LTR length *p* ≈ 0, tab), confirming that regulatory motif content and structural length vary systematically across LTR contexts. Solo LTRs consistently showed the lowest motif burden, motif density, and length, whereas LTRs adjacent to conserved internal domains (Full) were enriched for motifs and were longer (Figure 2B and supplemental Figures 1A and 1B).

For example, relative to solos, 3′-Full LTRs displayed a large increase in motif burden (Cliff’s δ = 0.486, FDR = 1.71×10^-19^), a medium increase in motif density (δ = 0.323, FDR = 2.63×10^-9^), and a medium increase in length (δ = 0.378, FDR = 2.67×10^-12^). Similar patterns held for 5′-Full LTRs (burden δ = 0.391, FDR = 1.60×10^-15^; density δ = 0.230, FDR = 3.19×10^-6^; length δ = 0.370, FDR = 6.23×10^-14^) and for intact proviruses retaining both LTRs (“Both-Full” category), where increases were strongest (burden δ = 0.560, FDR = 8.73×10^-189^; density δ = 0.364, FDR = 1.91×10^-80^; length δ = 0.483, FDR = 6.58×10^-141^).

Direct contrasts between conserved and degraded contexts reinforced this trend. 3′ LTRs flanking conserved internals carried more motifs and were slightly longer than 3′ LTRs adjacent to degraded regions (motif burden Cliff’s δ = 0.367, FDR = 1.38×10^-11^; motif density δ = 0.331, FDR = 1.59×10^-9^; length δ = 0.161, FDR = 3.5×10^-3^). Similar though somewhat smaller effects were observed for 5′ LTRs (burden δ = 0.310, FDR = 3.87×10^-10^; density δ = 0.253, FDR = 4.64×10^-7^; length δ = 0.219, FDR = 1.47×10^-5^). The largest Full–Deg differences appeared in conserved proviruses retaining both LTRs (“Both-Full”), where increases remained robust across all measures (burden δ = 0.409, FDR = 9.69×10^-84^; density δ = 0.283, FDR = 1.34×10^-40^; length δ = 0.298, FDR = 5.15×10^-45^).

As an internal symmetry check, 3′-Full vs 5′-Full differences were negligible across all measures (motif burden δ = 0.05, FDR = 0.46; motif density δ = 0.10, FDR = 0.19; length δ = −0.02, FDR = 0.77), indicating comparable regulatory preservation at both ends of conserved proviruses.

Finally, contrasts between degraded contexts and solo LTRs revealed only small or negligible effects, consistent with modest residual structure in degraded proviruses and extensive erosion in solos. For example, 3′-Deg LTRs carried only slightly more motifs than solos (motif burden δ = 0.151, FDR = 2.06×10^-102^; motif density δ = 0.008, FDR = 0.25) but were somewhat longer (length δ = 0.213, FDR = 1.16×10^-203^). Similar weak differences were seen for 5′-Deg (burden δ = 0.100, FDR = 1.74×10^-43^; density δ = –0.014, FDR = 0.067; length δ = 0.163, FDR = 6.53×10^-111^). Slightly stronger but still modest effects characterized Both-Deg proviruses, which retained both LTRs but degraded internals (burden δ = 0.187, FDR = 1.12×10^-96^; density δ = 0.082, FDR = 1.13×10^-19^; length δ = 0.196, FDR = 2.17×10^-106^). These results suggest that while degraded proviruses preserve marginally more sequence and regulatory content than isolated solos, both categories show advanced structural erosion relative to conserved loci.

Collectively, these data show that LTRs adjoining conserved internal coding potential are simultaneously longer and richer in TF motifs—in absolute burden and per-bp density—while solos and degraded contexts are shorter and motif-sparse. The co-variation of length, motif content, and internal domain integrity is consistent with coordinated selective maintenance of structural and regulatory features at a subset of HERV loci, providing plausible substrates for functional retention or co-option.

Consistent with this regulatory specialization, we observed that 5′ LTRs were approximately fivefold more likely than 3′ LTRs to overlap annotated transcription start sites (4.6 % vs 0.9 %; Supplemental Table 1). This enrichment aligns with the promoter function typically adopted by 5′ LTRs following integration, whereas 3′ LTRs—functioning as transcriptional terminators—are largely excluded from host TSSs. Although both LTRs originate as identical U3–R–U5 units, their divergent evolutionary fates have led to selective retention of promoter activity in 5′-like copies and polyadenylation features in 3′-like ones.

### U3–R–U5 Architecture Is Preferentially Preserved in Structurally Intact HERV Proviruses

To assess the structural integrity of HERV LTRs, we delineated the canonical retroviral subregions U3, R, and U5 across all elements using our motif- and geometry-aware annotation pipeline. This approach integrates promoter and polyadenylation signal models to infer transcriptional start and cleavage sites and reconstruct the full U3–R–U5 architecture, even in partially degraded LTRs. Benchmarking against shuffled LTR controls indicated that real genomic LTRs exhibit higher U3–R–U5 structural coherence than randomized sequences (Supplemental Results).

Each LTR was assigned a confidence label (“OK” or “LOW_CONF”) based on internal consistency between promoter and PAS calls, correct U3–R–U5 ordering, and the presence of auxiliary motifs such as PBS and PPT when applicable. Across all classes, roughly two-thirds of LTRs (66 %) were confidently assigned a complete U3–R–U5 architecture, confirming that vestiges of retroviral transcriptional organization remain broadly preserved in the human genome.

Proportions of confidently annotated (“OK”) LTRs were generally higher for loci flanking conserved internal domains than for degraded or solo elements. 3′-Full and Both-Full LTRs showed the highest annotation confidence (91 % and 92 %, respectively), followed by 5′-Full elements (79 %). In contrast, degraded proviruses displayed intermediate values (∼72–75 %), and solo LTRs were the least preserved (65 %) (Supplemental Table 2). Pairwise proportion tests confirmed that most Full vs Deg and all Full vs Solo contrasts were highly significant (BH-adjusted p < 0.001), with the exception of the 5′ Full vs 5′ Deg comparison. Absolute differences in OK fraction ranged from 0.07 to 0.27.

When comparing LTR sides, 3′ elements tended to exhibit higher annotation confidence than their 5′ counterparts within the same internal-domain context, an effect that was particularly pronounced for proviruses retaining conserved internal domains. This pattern is consistent with slightly faster structural erosion at the 5′ ends of proviruses, potentially reflecting asymmetrical selective constraints acting on promoter- versus terminator-associated regions. Notably, proviruses retaining both LTRs showed the highest overall preservation of U3–R–U5 structure (92 % OK when adjacent to conserved internal domains), indicating that when the full proviral architecture is maintained, both LTRs tend to remain jointly intact. Together, these observations support a model in which paired LTRs within complete proviruses exhibit coordinated structural maintenance, whereas isolated or solo LTRs are subject to more independent and variable erosion.

Focusing on the HERVH, HERVK, HERV17, HERVE, and HERVL subfamilies, clear differences emerged in the preservation of U3–R–U5 architecture (Fig. 1B). HERVH and HERVK displayed the highest proportions of well-preserved structures (95% and 92%, respectively), followed by HERV17 (89%), HERVE (76%), and HERVL (75%). Although overall preservation levels varied among families, a consistent asymmetry was observed within each: 5′ LTRs were more degraded than 3′ LTRs (Figure 2C), even in loci with well-preserved internal regions, a pattern that may reflect reduced promoter integrity at the 5′ end.

### Genomic Context Shapes the Internal Regulatory Architecture of LTRs

To investigate whether TFBMs are preferentially located at specific positions within LTR sequences, we performed positional density analysis across all motif matches identified by FIMO. For each LTR instance, motif match positions were scaled relative to LTR length and aligned in the 5′-3′ direction to compute kernel density estimates of motif distribution across different LTR classes.

When stratifying LTRs by type—5′ LTRs, 3′ LTRs, and solo LTRs—and by their distance to the nearest gene (categorized as at TSS, ≤1 kb, 1–10 kb, 10–100 kb, or >100 kb from TSS), we observed consistent patterns of positional enrichment. Notably, both 5′ LTRs and solo LTRs located directly at TSSs of known genes exhibited strong positional enrichment for several TFBMs (Fig 2C; e.g., DUX family motifs in 5′ and solo LTRs, FoxQ in 5′ LTRs, and ZBTB33 in solo LTRs), consistent with promoter-associated regulatory configurations. Interestingly enough, solo LTRs showed a marked accumulation of TFBMs near their 5′ end.

By contrast, 3′ LTRs displayed weaker and more diffuse positional patterns, especially when located at or near TSSs, with no consistent enrichment near either end of the LTR. This finding supports the hypothesis that 3′ LTRs—originally derived from the transcriptional terminus of proviruses—are under less selective pressure to preserve promoter-like regulatory elements compared to their 5′ counterparts.

Together, these results indicate that the internal regulatory architecture of LTRs is not uniformly distributed but instead reflects distinct positional organization of TFBMs. This bias is shaped by both structural context (solo vs. 5′ vs. 3′) and genomic positioning relative to host genes, pointing to functional specialization and selective retention of regulatory potential during LTR co-option.

### Solo LTRs at Gene Promoters Exhibit Enhanced Regulatory Architecture

A subset of solo LTRs overlaps the TSSs of annotated genes, placing them in a position to act as putative promoters (Table 1). To explore the regulatory potential of these elements, we focused on solo LTRs located at or near (<1 kb) TSSs and evaluated their TFBM content, enrichment profiles, and functional associations with nearby genes.

Analogously as above, we quantified the total number of TFBM matches per LTR, the motif density and the LTR’s length, and compared those solo LTRs overlapping TSSs (n = 5,138) to those located elsewhere in the genome (n = 532557). Across all three measures, at-TSS solos were significantly distinct from non-TSS solos (Kruskal–Wallis p ≈ 0). At-TSS elements exhibited a medium increase in motif burden (Cliff’s δ = 0.453, *FDR* ≈ 0), a small-to-medium increase in motif density (δ = 0.321, *FDR* ≈ 0), and a medium increase in LTR length (δ = 0.382, *FDR* ≈ 0). As exposed above, at-TSS solos also displayed sharp positional clustering of motifs toward the 5′ end of the LTR.

Consistent with these patterns, at-TSS solo LTRs also showed higher structural (U3-R-U5) annotation confidence: 80.6% of at-TSS solos were classified as “OK,” compared with 65.0% of solo LTRs located outside TSS regions. Together, these findings indicate that a subset of solo LTRs overlapping gene promoters may have been co-opted as regulatory modules, where both sequence length and motif enrichment contribute to their transcriptional activity.

### Solo LTR Promoters Are Enriched for Proliferative TF Motifs and Depleted of Neuronal Regulators

To investigate the regulatory potential of solo LTRs at TSS, we performed a differential motif enrichment analysis by comparing the frequency of TFBMs in solo LTRs located at TSS against a broader background of all LTR elements across the genome. Specifically, we considered motifs as present or absent per LTR, ensuring that repeated occurrences of the same motif within a single element did not inflate enrichment estimates. This allowed us to detect transcription factors whose motifs are significantly overrepresented—or depleted—in solo LTRs positioned at TSS, suggesting distinct regulatory configurations in promoter-proximal solo elements.

We found that a majority of TFBMs were significantly enriched in TSS-overlapping solo LTRs, with 595 out of 731 motifs exhibiting enrichment (odds ratio > 1, p-adjusted < 0.05). Among the most strongly enriched were motifs for transcription factors such as ZBTB14, HINFP, E2F1, TCFL5, KLF15, ZBTB33, E2F4, ZBED4, SPDEF, and E2F2. Many of these are associated with early developmental processes, cell cycle regulation, and with chromatin-opening roles—particularly members of the E2F family and zinc finger transcription factors like ZBTB33 and ZBTB14 (Du et al. 2025; Takebayashi-Suzuki et al. 2018; Pozner et al. 2016; Galán-Martínez et al. 2022). These findings support the hypothesis that co-opted solo LTRs often harbor regulatory architectures conducive to initiating or modulating transcription in developmental or proliferative contexts (Thompson et al. 2016; Ito et al. 2017).

However, the most striking insights arose from the small subset of depleted motifs—only 15 TFBMs were significantly underrepresented in TSS-overlapping solo LTRs compared with background, whereas the vast majority showed either enrichment or negligible effects. The top depleted motifs included CUX1, LHX3, HMGA1, PHOX2B, DLX2, DLX5, HOXB8, HOXB6, NEUROG1, and ZNF35, which include transcription factors involved in neural lineage specification, morphogenesis, and differentiation programs (Lu et al. 2023; Fedoseyeva et al. 2023; Glover 2001; Barretto et al. 2020; Ding et al. 1997; Dubreuil et al. 2002; Gandhi et al. 2020; Thaler et al. 2002; Cubelos et al. 2008), with ZNF35 showing preferential expression in neural progenitor and neural crest cell populations based on single-cell atlas data (Tomic et al. 2024).

To further assess the biological relevance of the depleted motifs, we performed GO enrichment analysis based on the transcription factors themselves. The analysis identified several significantly overrepresented biological processes (adjusted p-value < 0.05), predominantly related to neuronal differentiation and morphogenesis (cell projection organization, plasma membrane bounded cell projection organization, neuron projection development, cell morphogenesis involved in neuron differentiation). Several additional categories narrowly missed the significance threshold (adjusted p-value ≈ 0.1) but were functionally coherent, such as skeletal system morphogenesis, positive regulation of neuron differentiation, cell morphogenesis, and various cell projection morphogenesis terms (Figure 3A). Together, these results delineate a neuronal morphogenesis signature that contrasts with the profile of enriched motifs, which was dominated by regulators of early development and proliferation.

**Figure 3.**
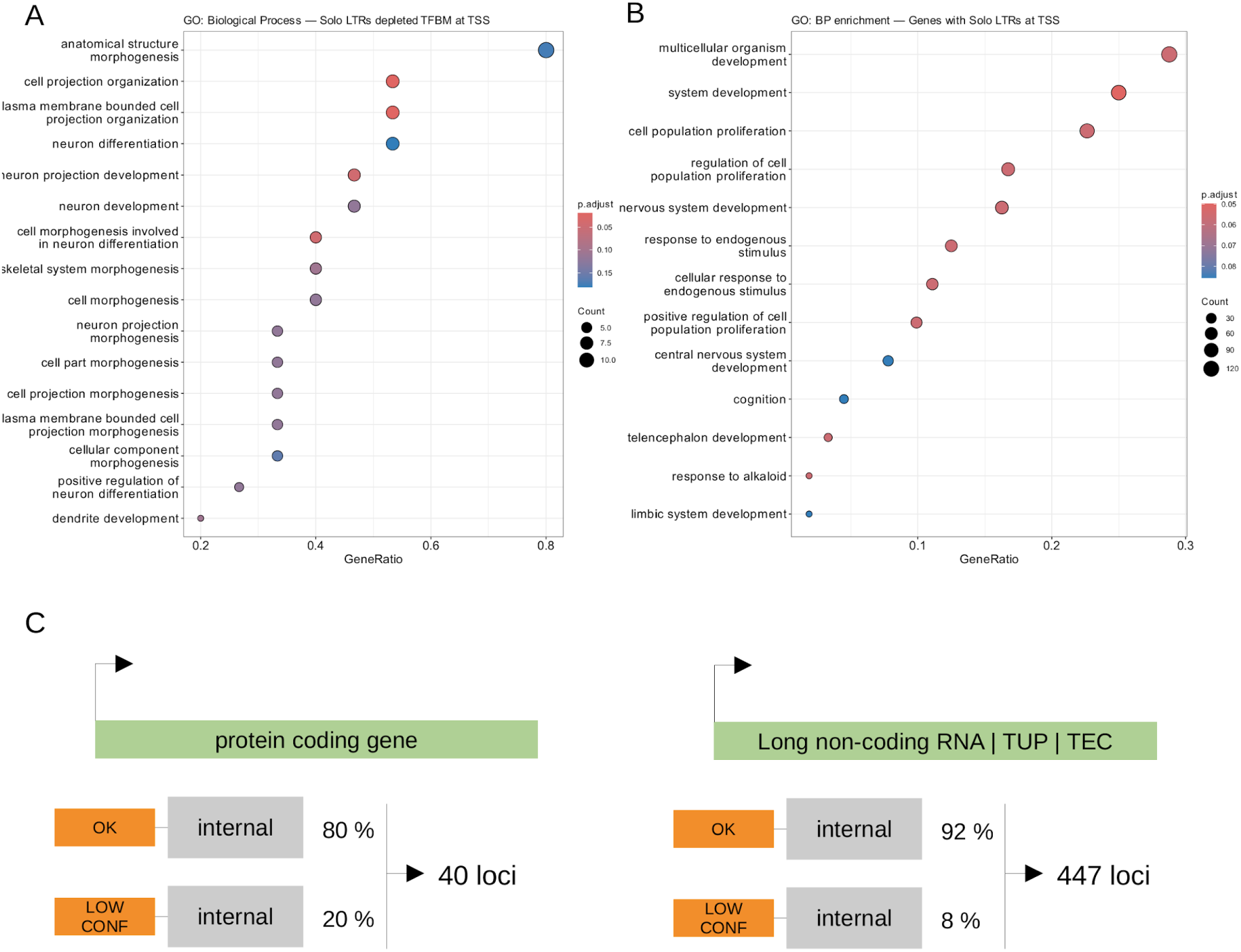
A) GO enrichment analysis of transcription factors whose binding motifs are present in solo LTRs overlapping the TSS of canonical genes. Many of the top enriched terms are associated with neuronal differentiation. B) GO enrichment analysis of genes with solo LTRs at their TSS. Numerous terms related to early development and morphogenesis emerge among the most significant categories. C) Summary of the 5′ LTR–at–TSS analysis. Hundreds of genes annotated as long non-coding RNAs appear to have HERV-derived promoter and internal regions segments.

Together, these findings indicate that solo LTRs co-opted as promoters selectively exclude motifs associated with neuronal differentiation, while favoring regulatory architectures that sustain or initiate transcriptional programs characteristic of early developmental and progenitor-like cellular states.

### Solo LTR Promoters Are Linked to Early Developmental and Proliferative Programs

To gain insight into the potential functional impact of co-opted solo LTRs acting as promoters, we performed GO enrichment analysis based on the nearest gene to each solo LTR overlapping a TSS.

The goal was to assess whether these loci, which exhibit significant TFBM enrichment, tend to regulate genes involved in specific biological processes as hypothesized above based on most strongly enriched motifs.

The GO enrichment analysis highlighted biological processes associated (adjusted p-value < 0.1) with developmental and proliferative programs (Figure 3B), including system development, multicellular organism development, nervous system development, central nervous system development, telencephalon development, limbic system development, and cognition, alongside regulation of cell population proliferation and related proliferation terms. These categories align with the motif-level findings, which featured TFBMs linked to developmental and cell-cycle regulators such as ZBTB33 and E2F1/2/4.

Taken together, these results indicate that solo LTRs overlapping TSSs preferentially associate with genes involved in developmental and proliferative programs, consistent with their enrichment for motifs linked to early regulatory factors. In contrast, motifs and GO terms related to differentiated or post-mitotic states were not overrepresented in this subset.

### lncRNA-Associated HERV Loci Retain Higher Promoter Integrity and Domain Conservation

To investigate whether HERVs contribute to the transcriptional landscape of uncharacterised loci, we focused on 5′ LTRs that overlap annotated TSSs. These LTRs were grouped into two main categories of LTR–gene associations. The first comprised 5′ LTRs located upstream of protein-coding genes, whereas the second encompassed transcripts annotated as non-protein-coding. Most instances in this second group (443 of 447) corresponded to long non-coding RNAs (lncRNAs), while a few were classified as transcribed_unprocessed_pseudogene (4 loci) or TEC (to be experimentally confirmed; 1 locus) (Supplemental Table 3).

Among 5′ LTRs overlapping protein-coding genes, approximately 80 % (32 loci) displayed well-defined U3–R–U5 promoter structures, but only about 10 % (4 loci: *AKR1C3*, *ATXN3*, *ERVW-1*, and *TFPI*) were linked to conserved internal domains. This indicates that, in protein-coding contexts, HERV-derived promoters can remain structurally intact even when downstream retroviral sequences have undergone extensive degeneration, suggesting selective retention of promoter activity after integration near host genes. In contrast, the lncRNA group showed both higher promoter quality (≈ 90 % OK; 409 loci) and a slightly greater frequency of conserved (≥ 0.95 HMM model coverage) internals (≈ 12 %; 55 loci) (Figure 3C). Among these, protease (AP) domains were the most frequently conserved (found in 45 loci), followed by RNase H (16), reverse transcriptase (10), dUTPase (9), integrase (8), envelope (8), and GAG (7). In this group, catalytic pol-derived domains—particularly AP and RNase H—were more commonly preserved than structural (GAG) or envelope elements.

Taken together, these observations show that 5′ LTRs associated with lncRNA-annotated loci exhibit both higher promoter quality and a higher frequency of conserved internal domains than those linked to protein-coding genes, indicating a distinct structural profile for this group.

## Discussion

Our systematic annotation of HERV LTRs, integrated with our previous domain-level analysis of internal regions, reveals that these retroviral remnants are far from uniform genomic fossils. Instead, they preserve distinct layers of structural and regulatory organization that reflect both their evolutionary history and potential functional physiological current relevance, in health and disease. By jointly examining motif content, structural context, and U3–R–U5 architecture, we observe a consistent hierarchy of preservation across LTR classes. Notably, a subset of LTRs—particularly solo copies positioned at transcription start sites—retain dense, non-random clusters of promoter-associated motifs suggesting sustained evolutionary constraint at these loci. The non-random nature of these architectures was further supported by benchmarking the U3–R–U5 model against shuffled LTR controls, which exhibited substantially reduced structural coherence (Supplemental Results). These findings support the long-standing view that LTRs are frequently co-opted as regulatory elements (Thompson et al. 2016; Ito et al. 2017; Chuong et al. 2016; Frank and Feschotte 2017) and extend it by demonstrating that promoter-like features are selectively maintained in specific structural configurations, rather than being distributed randomly among degraded copies.

A central observation is that LTRs flanking internal HERV regions with well-preserved gag, pol, or env domains exhibit higher motif burden, higher motif density, and longer sequence length compared with solo LTRs. Direct contrasts with degraded proviruses reinforce this pattern, indicating a gradient in which LTRs adjoining conserved internal coding potential retain the greatest regulatory capacity, degraded contexts show intermediate values, and solos are the most eroded. This association between preserved internal domains and retained regulatory potential suggests that structural and regulatory features may have been maintained together at these loci, possibly reflecting evolutionary constraint or recurrent co-option rather than simple neutral decay (Frank and Feschotte 2017). Such co-maintenance is compatible with well-characterized examples of coordinated host recruitment, as in the case of syncytins, where both the envelope gene and its LTR promoter were exapted for placental development (Cornelis et al. 2015). Alternatively, regulatory sequences may have been preserved independently of coding potential, a process exemplified by MER41 LTRs, which were co-opted as interferon-inducible enhancers controlling immune genes (Thompson et al. 2016).

Our U3–R–U5 segmentation analysis provides a complementary structural explanation for this pattern, revealing that the transcriptional architecture itself is retained preferentially at the same loci that show elevated motif burden and preserved internal domains. LTRs that flank internal coding regions consistently retain more complete U3–R–U5 architecture than degraded or solo copies, reinforcing the structural–regulatory gradient observed in our motif and length analyses. In intact proviral contexts (those still keeping the structure LTR-internal-LTR), both the upstream U3/R promoter module and the downstream U5/PAS termination module tend to remain preserved, whereas degraded and solo LTRs show progressive loss of these features. This coordinated maintenance of paired promoter and termination signals suggests that certain proviruses retained not only coding potential but also coherent transcriptional directionality, in contrast to the more fragmented architecture seen in the majority of copies.

A consistent asymmetry emerged across families: 3′ LTRs were generally more intact than 5′ LTRs, implying that termination-associated signals are more resistant to erosion than promoter modules after endogenization. This pattern aligns with the asymmetric selective pressures acting on proviral LTRs: 5′ LTRs carry the promoter responsible for initiating viral transcription and are therefore prime targets for silencing and mutational erosion, whereas 3′ LTRs lack promoter activity and experience comparatively weaker selection, allowing their U5/PAS modules to remain structurally intact for longer periods. Subfamily analyses further support this view: younger families such as HERVH and HERVK show markedly higher retention of full U3–R–U5 units than older families such as HERVE or HERVL, mirroring known evolutionary age effects.

These findings extend our earlier motif-based observations by suggesting that the regulatory potential of LTRs resides not only in isolated binding sites but in the preservation of the entire promoter–initiation–termination module. This structural coherence is concentrated in the same subset of loci that retain internal coding domains, arguing in favour of these proviruses maintaining both their regulatory and structural components for longer periods, thereby increasing their likelihood of being exapted as functional promoters or terminators in the host genome. In contrast, the majority of solo or degraded LTRs show partial or stochastic retention of subregions, consistent with widespread neutral erosion, and distinguishing a minority of structurally coherent loci that may have exerted disproportionate evolutionary influence (Thompson et al. 2016).

At the regulatory level, solo LTRs overlapping TSSs exhibit widespread motif enrichment, with most tested motifs overrepresented compared to background. Among these, motifs for early developmental and proliferative regulators (e.g. E2Fs (Du et al. 2025), ZBTB14/33 (Takebayashi-Suzuki et al. 2018; Pozner et al. 2016), TCFL5 (Galán-Martínez et al. 2022)) are among the most strongly enriched, while a small subset of motifs linked to neuronal differentiation (e.g. NEUROG1 (Lu et al. 2023), HOXB6 (Fedoseyeva et al. 2023; Glover 2001), DLX2 (Barretto et al. 2020; Ding et al. 1997)) are consistently depleted. This asymmetry is also reflected in gene ontology, where nearby genes are enriched for developmental and progenitor functions but depleted for differentiation categories. Enrichment suggests active recruitment of pioneer and developmental regulators, while depletion is equally informative: it implies a systematic exclusion of architectures incompatible with progenitor-like states. Such pruning likely reflects host-mediated selection after endogenization: differentiation-associated motifs would drive LTR activity in somatic or late developmental contexts where expression is deleterious to the host and offers no benefit to vertically inherited ERVs, whereas motifs compatible with the permissive regulatory environment of early embryonic stages are more readily tolerated (Efroni et al. 2008).

These observations resonate with the evolutionary framework outlined by Thompson et al. (2016) (Thompson et al. 2016): ERVs have likely been under selection to express in germline and early embryonic contexts, taking advantage of epigenetic reprogramming windows in preimplantation embryos and placenta that permit ERV transcription despite KRAB-ZFP/KAP1-mediated repression (Thompson et al. 2016). LTRs enriched in transcription-factor binding sites can autonomously recruit host TFs, increasing transcriptional output and the probability of vertical transmission (Thompson et al. 2016). Our data suggest that the same promoter architectures that likely enabled viral propagation are those that persisted in the host genome, later co-opted as developmental gene promoters. The enrichment of early developmental regulators and depletion of differentiation factors in TSS-overlapping solo LTRs thus represent an evolutionary “imprint” of ERV biology.

Our findings also intersect with the emerging literature on transposable elements and lncRNA biology. Transposable elements account for a substantial fraction of lncRNA sequence, with 75–83% of human lncRNAs containing at least one TE and LTRs frequently acting as promoters that initiate their transcription (Fort et al. 2021). We extend this view by showing that a subset of transcripts annotated as lncRNAs in current catalogs likely represent transcriptionally active HERV loci with preserved internal domains and residual coding capacity. The co-occurrence of promoter-like LTRs, intact U3–R–U5 promoter modules, and preserved retroviral domains argues that these are candidate mis-annotations, blurring the boundary between noncoding and coding transcription. If such loci produce peptides or proteins, they could participate in processes ranging from placental development—classically exemplified by syncytins (Cornelis et al. 2015)—to oncogenesis (Göke and Ng 2016) and immune modulation (Chuong et al. 2016). Illustrative precedents already exist, as LTR-driven lncRNAs such as linc-RoR, HPAT5, and SAMMSON have been shown to influence pluripotency, early embryogenesis, and cancer (Göke and Ng 2016). By systematically highlighting these candidates, our study provides a roadmap for experimental re-evaluation of the lncRNA landscape and the hidden coding potential of endogenous retroviruses.

More broadly, these results underscore how co-opted LTRs have provided a recurrent regulatory substrate for rewiring transcriptional programs in pluripotency and immunity (Thompson et al. 2016). The systematic biases we observe—favoring developmental regulators while excluding differentiation motifs, retaining complete U3–R–U5 regulatory modules at select loci, and coupling regulatory integrity with preserved internal domains—may suggest that LTR-driven promoters have been preferentially retained for roles in progenitor states.

Our analyses are based primarily on motif content, positional architecture, and genomic context, which do not by themselves establish regulatory activity. Functional engagement depends on chromatin state and transcription factor availability in specific cell types. LTR overlap with internal exons or isoforms was not exhaustively resolved, raising the possibility of unrecognized impacts on transcript isoform diversity. Moreover, while our classification distinguishes solo, 5′, and 3′ LTRs, many proviral insertions in the genome are degraded in more complex patterns—such as tandem LTRs or nested insertions—that this framework does not fully capture. Finally, we did not investigate sequence conservation across species; we considered that many co-opted LTRs are relatively young and lineage-specific (Jacques et al. 2013), thus they would not necessarily display cross-species conservation signals, even if they are functional in humans.

To help address these challenges, we developed HERVarium, an interactive platform for exploring HERV structure (internal domains and LTRs) and regulatory potential. Beyond the systematic annotation of LTRs, internal domains, and U3–R–U5 architecture, the interface integrates chromatin accessibility and expression tracks (e.g., GTEx), enabling visualization of HERV annotations in the context of tissue- and cell-type–specific activity. This integration facilitates hypothesis generation by allowing researchers to rapidly assess whether candidate LTRs or proviral loci coincide with open chromatin or transcriptional activity. In its current version, HERVarium is distributed for local installation via the project’s GitHub repository, ensuring transparency and reproducibility. A public web-accessible release is planned for the time of publication to enable direct online exploration. Although we have not systematically investigated these additional data layers here, HERVarium provides a framework for cross-referencing retroviral annotations with functional genomic datasets, supporting both hypothesis-driven follow-up studies and community-based exploration.

Building on this foundation, future work will integrate high-resolution epigenomic and transcriptomic datasets (e.g., ATAC-seq, ChIP-seq, single-cell RNA-seq, ERV-seq) into HERVarium to more precisely identify LTRs with bona fide regulatory activity across developmental, aging and disease contexts. CRISPR-based perturbations, as recommended by Thompson et al. (2016) (Thompson et al. 2016), now offer a feasible strategy to test the functional significance of candidate LTR-driven promoters and enhancers in vivo. Comparative analyses across primates, where many insertions are species-specific, will help clarify how lineage-specific insertions contributed to regulatory evolution.

Finally, expanding HERVarium to additional mammalian genomes and coupling it with evolutionary conservation metrics will help reveal whether the co-retention of U3–R–U5 modules and internal coding capacity is a general property of ERVs or a trait accentuated in humans.

## Methods

### Identification and preprocessing of LTRs

Genomic coordinates of long terminal repeats (LTRs) were obtained from the human reference genome (GRCh38) using RepeatMasker (Tarailo-Graovac and Chen 2009) v4.1.8, with NCBI/RMBLAST (v2.14.1+) as the search engine and the Dfam 3.9 library (Storer et al. 2021). Only entries classified as LTR/ERV were retained, while internal regions annotated as “–int” or “_I” were excluded. To reduce redundancy and approximate transcript-like proviral configurations, nearby LTRs from the same subfamily and strand were merged when separated by ≤100 bp, resulting in a non-redundant set of LTR intervals.

### Sequence extraction and stratification

Merged intervals were used to extract strand-specific genomic sequences from GRCh38 with Biopython and pyfaidx (v0.8.1.4). The resulting multi-FASTA file was partitioned by subfamily, generating separate FASTA files to enable motif scanning in parallel.

### Motif scanning

LTR sequences were scanned for TFBMs using FIMO (Grant et al. 2011) (MEME Suite (Bailey et al. 2015) v5.5.8). Background models were generated with fasta-get-markov from the complete set of LTR sequences, and motif scanning was performed separately for each subfamily FASTA file with a p-value threshold of 1×10^-4^. To limit memory usage, the maximum number of stored hits was set to one million per run.

### Aggregation and coordinate mapping

Subfamily-specific FIMO results were merged into a single table, with each motif hit annotated by its originating LTR sequence and subfamily. Motif positions were then mapped back to genomic coordinates, producing both a sorted TSV and a BED file with per-hit coordinates suitable for downstream visualization (Figure 4).

**Figure 4.**
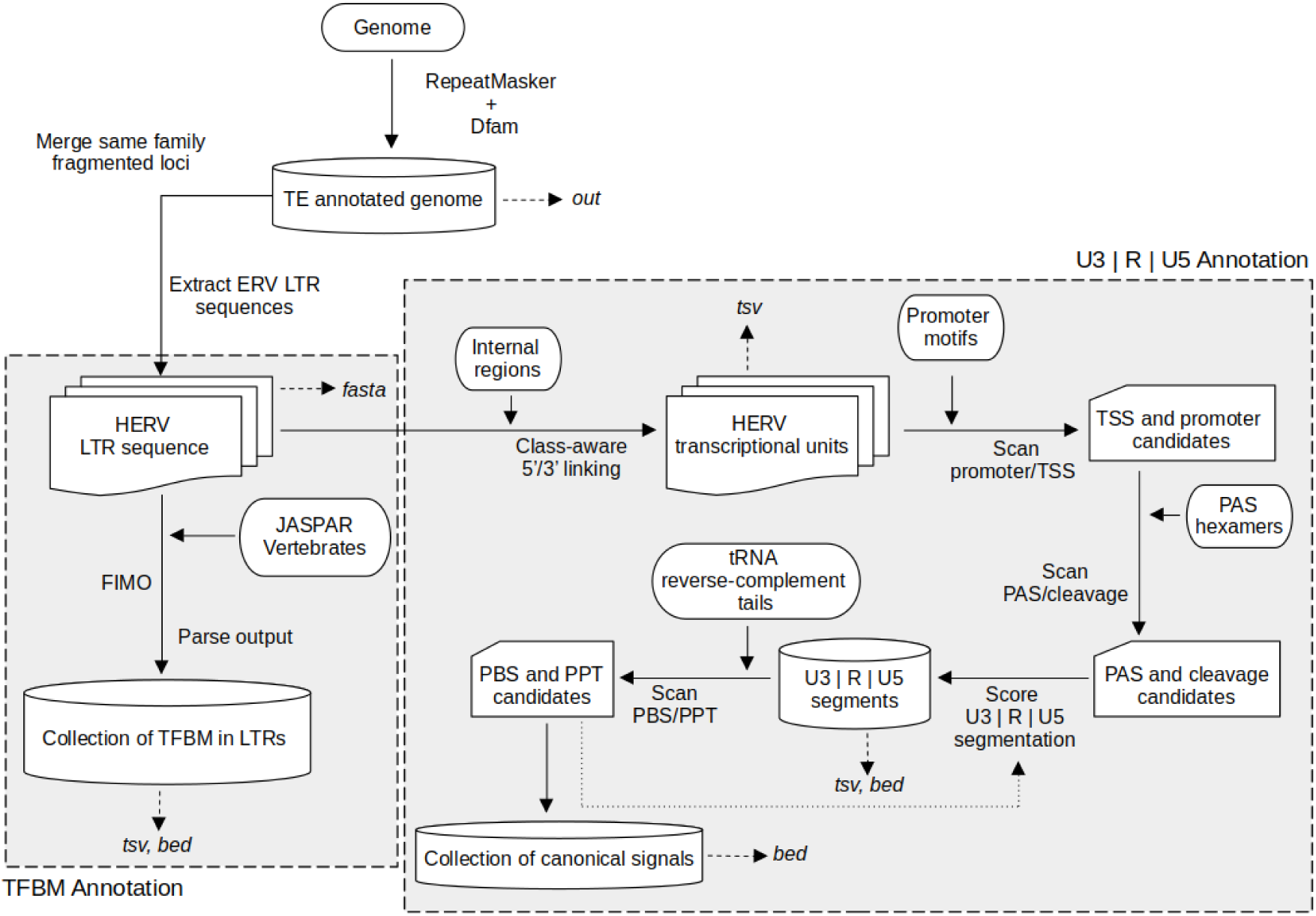
Overview of the annotation pipeline. The workflow begins with RepeatMasker output and proceeds through two complementary analyses of LTR sequences: (i) TFBM annotation using FIMO with the JASPAR database, and (ii) U3/R/U5 segmentation together with PBS and PPT detection using a custom Python script informed by promoter motif definitions, PAS hexamers, and reverse-complement tRNA tails. While the main output is a TSV file containing domain coordinates and completeness scores, several intermediate and supporting files are also generated, including FASTA, TSV, BED, and TBL files, as well as InterProScan and Phobius output files providing additional information for each detected domain.

### Integration with internal regions and domain annotations

Motif coordinates were integrated with our previously generated domain-level HERV annotation (Montserrat-Ayuso et al. 2026), which includes GyDB/InterProScan–derived protein-domain calls (*env*, *gag*, *pol*, accessory) and conservation tiers defined from HMM profile coverage. LTRs were linked to the nearest internal region within ±200 bp on the same strand and ERV class, and classified as 5′, 3′, both, or solo relative to proviral orientation. This yielded a combined dataset linking LTRs, internal regions, and domain conservation status.

### U3–R–U5 annotation and PBS/PPT detection

The canonical retroviral subregions U3, R, and U5 were annotated across all LTR sequences using a motif-based and geometry-aware workflow that detects degenerate promoter and polyadenylation elements and evaluates their consistency with expected retroviral organisation. Promoter-associated motifs (TATA box, Initiator [Inr], downstream elements such as DPE and MTE, BREu/BREd, SP1, and related GC-boxes) were detected using consensus sequence patterns and spacing rules derived from established core-promoter models (Butler and Kadonaga 2002; Juven-Gershon et al. 2008; Kugel and Goodrich 2017), reflecting the highly degenerate nature of core-promoter elements in endogenous retroviruses. Candidate polyadenylation signals in U5 were identified using canonical and variant PAS hexamers (Beaudoing et al. 2000), together with their associated CA cleavage dinucleotide and downstream GU-rich elements. Putative transcription start sites and PAS/cleavage sites were evaluated using soft, biology-informed geometry heuristics that weakly favoured canonical U3, R, and U5 lengths (>70 bp for U3, 60–350 bp for R, and ∼100 bp for U5), consistent with retroviral LTR organization (Coffin et al. 1997), from which the most internally consistent U3–R–U5 segmentation was selected for each LTR.

For 5′ and 3′ LTRs, flanking regions were scanned for primer binding sites (PBS) and polypurine tracts (PPT), respectively. PBS motifs were matched against reverse-complement 3′ tRNA-tail sequences derived from mature human tRNAs obtained from GtRNAdb (https://gtrnadb.org/genomes/eukaryota/Hsapi38/hg38-mature-tRNAs.fa) (Chan and Lowe 2009, 2016), allowing limited mismatches to accommodate ERV-specific divergence. PPT motifs were detected as purine-rich tracts characteristic of retroviral replication intermediates (Coffin et al. 1997).

Each LTR was assigned a confidence label (OK or LOW_CONF) based on the internal consistency of detected motifs and spacing constraints, rather than on functional interpretation. Additional methodological benchmarking using randomized LTR controls is described in Supplemental Methods.

The resulting annotations were summarised as BED and TSV tables reporting genomic coordinates, motif evidence, and confidence metrics. Detailed algorithmic parameters are found in Supplemental Methods, and the full implementation is provided in the accompanying code repository (https://github.com/funcgen/herv-regulatory-map) (Figure 4).

### Motif burden and density

FIMO-derived motif hits were counted per LTR to quantify TFBM burden. Motif density was defined as the number of TFBM per base pair of LTR sequence. Counts were compared across LTR categories defined by the conservation status of adjacent internal regions (e.g., LTRs adjacent to intact vs. degraded domains, or solo LTRs). Domains were considered intact when the maximum HMM profile coverage exceeded 95%, reflecting near-complete conservation of the corresponding domain. Motif burden distributions were visualized as boxplots, with extreme outliers truncated at the 99th percentile for visualization purposes.

### Transcription start site annotation

Distances from each LTR to the nearest transcription start site (TSS) were computed using GENCODE (Mudge et al. 2025) v48 annotations in a strand-aware manner. Distances were binned into categories (at TSS, ≤1 kb, 1–10 kb, 10–100 kb, >100 kb), and each LTR was annotated with its nearest gene. Motif burden was analyzed as a function of TSS proximity, with particular emphasis on solo LTRs overlapping TSSs.

### Motif positional analysis

For each motif hit, the relative position within the LTR was calculated by normalizing the midpoint of the match to LTR length (scaled 0–1). Kernel density estimation (KDE) was then performed on relative positions, stratified by LTR class (5′, 3′, solo) and TSS proximity. Only strata—defined as motif × LTR class × TSS bin combinations—with at least 100 motif hits were retained for analysis. KDE curves were smoothed with a moving average and complemented with histogram-based estimates. Positional peaks were identified where the smoothed density exceeded both neighboring values, and peak sharpness was defined as the ratio between peak density and the mean density within each stratum.

### Motif and functional enrichment analyses

#### TFBM overrepresentation at putative co-opted promoters

We tested for overrepresentation of TFBMs in solo LTRs overlapping a TSS (foreground) relative to all other LTRs (background). FIMO hits were collapsed to unique LTR×motif pairs to avoid within-element multiplicity. For each motif, a 2×2 table (foreground/background × with/without motif) was evaluated by Fisher’s exact test, with Benjamini–Hochberg correction across motifs. Motifs were considered enriched if OR > 1 and FDR < 0.05. Results are shown as volcano plots using capped log2(OR) to stabilize extreme estimates.

### Gene Ontology (GO) enrichment—two complementary views

i. **TF-centric GO:** significantly enriched (and, where applicable, depleted) motif IDs were mapped to TF symbols and tested with clusterProfiler::enrichGO(Yu et al. 2012) (OrgDb = org.Hs.eg.db, ontology = BP, BH adjustment; reporting thresholds padj < 0.1, q < 0.2). The universe comprised all TFs represented among tested motifs.
ii. **Gene-centric GO:** independently, we performed GO on nearest genes to solo LTRs at TSS (study set) versus those within ≤1 kb of a TSS (comparison), using the union of both as the background.

### Output tables and reproducibility

All intermediate and final results were written to disk as TSV files, including: (i) LTR classification with TSS annotation, (ii) internal region + domain summaries, (iii) motif positional KDE and histogram tables, and (iv) motif enrichment results. The analysis workflow of FIMO results was implemented in R (v4.3.3) using data.table (Barrett et al. 2025), GenomicRanges (Lawrence et al. 2013), ggplot2 (Wickham 2016), clusterProfiler (Yu et al. 2012), and enrichplot.

### Interactive web application (HERVarium)

To enable rapid exploration of the HERV structural and regulatory annotations, we developed HERVarium, an interactive web application implemented in Python using Dash (v3.2.0) and the Bootstrap front-end framework for layout and styling. The interface offers real-time filtering, interactive locus-level visualization, and one-click export of filtered annotations.

### Precomputation of annotation tables

All annotation layers—LTR classifications, LTR–gene distance and TSS bins, motif burden, conserved internal domains, and U3–R–U5 features—were collated into a set of precomputed tables stored in Parquet format for efficient server-side querying. Identifiers were harmonized across datasets, normalized coordinate-based names, merged LTR and internal-region metadata, and integrated motif, PBS/PPT, and promoter/PAS signal annotations. Summary metadata (value ranges, categorical levels, feature lists) were exported as JSON to configure dropdowns and filters dynamically.

### Genome browser integration

A genome browser was embedded in the application using IGV.js (via dash-bio). All tracks used by the browser were generated from the processed BED files using a standardized BED to bigBed pipeline. Dedicated scripts ensured consistent formatting (BED6 compliance, simplified locus names, integerized scores) and reproducible bigBed creation using chromosome sizes derived from the GRCh38 primary assembly. Tracks included: (i) internal HERV regions and domain calls; (ii) LTRs; (iii) U3/R/U5 segments, PBS/PPT annotations, and promoter/PAS signals; (iv) genome-wide TFBM hits from FIMO; (v) GENCODE v48 gene models converted to bigBed using a custom pipeline and (vi) optional external datasets consisting of ENCODE DNase I hypersensitivity profiles and GTEx v8 tissue RNA-seq coverage (bigWig). All tracks are rendered on demand and synchronized with table selections.

### Backend and filtering logic

Annotation tables for LTRs and internal HERV loci are loaded at startup, while U3/R/U5 features are queried on demand using a persistent DuckDB in-memory connection. The application supports filtering by LTR subfamily, LTR class (solo, 5′, 3′), linked internal subfamily, motif count, genomic length, domain presence and HMM coverage, U3/R/U5 feature class, segment confidence, and TSS distance. Filters persist across sessions and update both the annotation tables and the genomic locus displayed in IGV. All tables support sorting, pagination, and export to CSV.

### Deployment and availability

HERVarium is packaged as a standalone Dash application and can be deployed on any standard Python environment. The full source code, including preparation scripts and the web application, is available at GitHub (https://github.com/funcgen/HERVarium) and Zenodo (https://doi.org/10.5281/zenodo.18551738).

## Supporting information

Supplemental Results

## Data access

The human genome assembly GRCh38 can be downloaded from the GENCODE website: https://ftp.ebi.ac.uk/pub/databases/gencode/Gencode_human/release_49/GRCh38.primary_assembly.genome.fa.gz.

The transcription-factor binding motifs were obtained from JASPAR 2024, specifically the CORE vertebrates non-redundant position frequency matrices (PFMs).

The Dfam database partitions used for RepeatMasker were downloaded from: https://www.dfam.org/releases/current/families/FamDB/dfam39_full.0.h5.gz (root), and https://www.dfam.org/releases/current/families/FamDB/dfam39_full.7.h5.gz (Mammalia).

The complete HERV LTR annotation datasets—including the RepeatMasker output files used for this analysis, BED and TSV files, U3–R–U5 segment definitions, PBS/PPT annotations, motif-scan outputs, and the supporting tRNAs, promoter, and PAS-hexamer definition files—is publicly available via Zenodo at DOI: https://doi.org/10.5281/zenodo.18554805.

## Code availability

The pipeline used to identify open reading frames, annotate retroviral domains and analyze the results is available as a collection of Python, Bash, and R scripts at https://github.com/funcgen/herv-domain-map.git.

## Acknowledgements

We acknowledge the use of artificial intelligence tools (ChatGPT, OpenAI) to assist in code drafting and language editing.

Author contributions: T.M-A and A.E-C conceived the project. T.M-A performed the bioinformatic analyses. T.M-A and A.E-C wrote the manuscript. A.E-C supervised the project. A.P. contributed to scientific discussions and provided critical feedback on the manuscript.

## Funding

This publication and all its results are supported by the AGAUR-FI predoctoral grant program (2025 FI-1 00642) Joan Oró, from the Secretariat for Universities and Research of the Department of Research and Universities of the Government of Catalonia, and by the European Social Fund Plus. AP has received funding from the Secretariat for Universities and Research of the Ministry of Business and Knowledge of the Government of Catalonia (2021SGR00899), and the Instituto de Salud Carlos III (ISCIII) (FIS PI23/00835), ‘Fondo Europeo de Desarrollo Regional (FEDER), Unión Europea, una manera de hacer Europa’. Institutional support to CNAG was from the Spanish Ministry of Science, Innovation and Universities, Fondo de Investigaciones Sanitarias cofunded with ERDF funds (PI19/01772), the 2014–2020 Smart Growth Operating Program, and the Generalitat de Catalunya through the Departament de Recerca i Universitats and Departament de Salut.

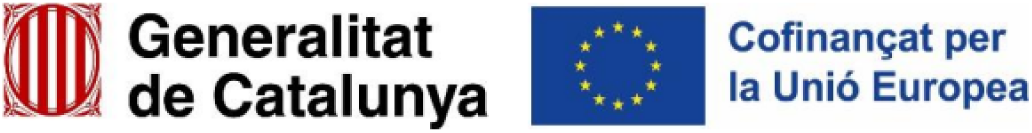

## Supplemental Tables

Supplemental Table 1: Genomic distribution of HERV LTRs relative to annotated transcription start sites (TSS). Percentages indicate the fraction of 5′ LTRs, 3′ LTRs, and solo LTRs located at the TSS or within increasing distance bins from the nearest annotated TSS.

Supplemental Table 2: Annotation confidence of HERV LTRs across structural contexts. Shown are the proportions of LTRs with non-reconstructed versus reconstructed U3–R–U5 architecture, stratified by LTR position and internal domain conservation status.

Supplemental Table 3: Catalogue of 5′ HERV LTRs overlapping annotated transcription start sites (TSS), classified by their association with protein-coding genes or non-protein-coding transcripts. Non-coding associations include long non-coding RNAs (lncRNAs), transcribed unprocessed pseudogenes, and TEC (to be experimentally confirmed) annotations.

