## Supplemental Methods for "Regulatory Features and Functional Specialization of Human Endogenous Retroviral LTRs: A Genome-Wide Annotation and Analysis via HERVarium"

### Annotation of U3, R, and U5 Regions in HERV LTRs

We annotated the canonical U3, R, and U5 subregions in all LTR sequences using a rule-based pipeline that integrates position-aware motif detection, using weighted motif evidence and spacing-aware scoring, and using empirically informed length preferences for U3, R, and U5 (Butler and Kadonaga 2002; Juven-Gershon et al. 2008; Kugel and Goodrich 2017; Coffin et al. 1997).

#### Input preparation and strand normalization

All LTR sequences were extracted in transcription (+) orientation, regardless of genomic strand, ensuring that promoter elements (upstream), the TSS (boundary of U3-R), and polyadenylation signals (downstream; R-U5) were evaluated in a consistent coordinate space. Flanking genomic windows (±200 bp by default) were also retrieved and normalized to the + orientation for PBS/PPT detection.

LTRs were labeled as 5′ and/or 3′ if they occurred within 200 bp (edge-to-edge, strand-aware) of an internal ERV region of the same ClassFamily; LTRs assigned at least once as both 5′ and 3′ across all internal regions were labeled ‘both’ and were scanned for both PBS and PPT signals.

#### Promoter analysis and TSS identification (U3 region)

#### Motif discovery: strict and permissive Initiator (Inr)

The pipeline first identifies candidate transcription start sites (TSS) using two Initiator models: Strict Inr: the canonical YYANWYY motif, with the TSS anchored at the conserved ‘A’ position. Permissive mammalian Inr (YR Inr): a relaxed dinucleotide Y–R motif, with the TSS anchored at the second base; these candidates carry a lower base weight. Candidate TSSs supported by both models at the same genomic position are deduplicated, preferentially retaining strict Inr-supported sites.

In LTRs lacking any Inr signal, a fallback strategy generates provisional TSS candidates from TATA box positions, placing the TSS at a fixed offset downstream.

**Scoring of promoter context**

Each candidate TSS is assigned a composite score integrating the presence and spacing of core promoter elements, according to table 1 (Butler and Kadonaga 2002; Juven-Gershon et al. 2008; Kugel and Goodrich 2017).

Table 1. Core promoter motifs, their expected positions relative to the TSS, and their contributions to the promoter score.

| **Motif** | **Expected position relative to TSS** | **Contribution** |
|---|---|---|
| TATA box | −30 ± 20 bp upstream | strongest positive weight (distance-scaled) |
| BREu/BREd | immediately flanking TATA | moderate weight (TATA-dependent) |
| DPE | +28 to +32 | strong downstream element, particularly informative for TATA-less promoters |
| MTE | +18 to +27 | auxiliary downstream promoter element |
| DCE (SI, SII, SIII) | +6–11, +16–21, +30–34 | small cumulative boosts |
| SP1 / GC-box | −150 to +10 | moderate activator weight |
| XCPE1 | −8 to +2 | very small bonus for TATA-less promoters |
| GC% bonus (optional) | −150 to +50 | applied only to TATA-less promoters in high-GC context |

The TATA contribution declines linearly as its spacing deviates from the ideal −30 bp. Downstream elements (DPE, MTE, DCE) and auxiliary motifs (SP1, XCPE1) provide additive support, with BRE elements evaluated only in the presence of a nearby TATA box. A mild penalty is applied to candidate TSSs yielding very short U3 regions (<60 bp), with reduced impact for promoters supported by strong motif evidence. An optional GC-content bonus can be applied to TATA-less promoters in high-GC contexts.

**TSS candidate list**

Each Inr-based candidate is represented as tss: <pos>, score: <float>, evid: <motif/evidence string>, has_tata: True/False. If no Inr-based candidates are found, a TATA-only fallback generates provisional TSS candidates by placing the TSS 30 bp downstream of each detected TATA motif (relative to the TATA match end).

**Polyadenylation signal and cleavage detection (U5 region)**

PAS detection proceeds in two stages. First, a canonical scan examines the 3′ portion of the LTR (≈280 bp) for PAS hexamers drawn from a curated list of canonical and variant signals. A candidate defines a cleavage site if a CA dinucleotide is detected 10–30 nt downstream; the presence of a GU-rich downstream element (DSE) increases confidence. The scan proceeds from 3′ to 5′, preferentially selecting the most downstream PAS with a valid cleavage site.

If the canonical scan yields no suitable cleavage site, a sensitive scan is applied to a wider region (≈360 bp). This second pass accepts canonical PAS hexamers or A-rich 6-mers (≥4 A), requiring a downstream CA and a GU-rich element for A-rich motifs, and enforces biologically plausible U5 lengths (hard minimum 40 bp; soft maximum 320 bp). Candidates are scored based on PAS type, DSE support, and U5 geometry.

After TSS selection, PAS detection is repeated on the sequence downstream of the TSS. If no valid PAS is found in this region, a previously detected PAS is retained only if it lies downstream of the selected TSS; otherwise, PAS is treated as absent. The final PAS record stores the PAS position, cleavage site, score, and supporting evidence.

**Coupled selection of U3 - R - U5 boundaries**

**Coupled evaluation of promoter and PAS candidates**

For each promoter candidate, the pipeline evaluates its compatibility with the PAS. R length (distance from TSS to PAS cleavage) is scored relative to an expected window (≈60–350 bp) using a smooth, bell-shaped tolerance function. U5 length contributes an additional soft preference term (ideal ≈100 bp). Very short U3 regions incur mild penalties, reduced when promoter evidence is strong. The optimal TSS is selected by maximizing: joint_score = promoter_score + R_prior + (0.5 × U5_prior) − U3_penalty. If PAS evidence is absent or incompatible with promoter geometry, the highest-scoring standalone promoter candidate is selected.

**Defining U3, R, and U5**

Final segmentation defines U3 as the interval from the LTR start up to (but not including) the TSS; R from the TSS up to (but not including) the PAS cleavage site; and U5 from the cleavage site to the end of the LTR. Segments are mapped back to genomic coordinates in a strand-aware manner. Segments are mapped back to genomic coordinates in a strand-aware manner. LTRs lacking valid TSS or PAS signals, or exhibiting invalid geometry, are labeled LOW_CONF.

**Annotation of PBS and PPT in flanking regions**

**Primer Binding Site (PBS, downstream of 5′ LTRs)**

Downstream flanks of 5′ or ambiguous LTRs were scanned for tRNA-derived primer binding sites. Reverse-complemented 3′ tRNA sequences from GtRNAdb (Chan and Lowe 2009, 2016) were deduplicated, and variable-length suffixes (12–20 nt) were matched using fuzzy Hamming-distance tolerances (~15% mismatches). Matches were scored by alignment length and mismatch count, with a boundary-proximity penalty favoring sites closest to the LTR edge. The highest-scoring match was reported as the PBS signal.

**Polypurine Tract (PPT, upstream of 3′ LTRs)**

Flanks upstream of 3′ or ambiguous LTRs were scanned for purine-rich tracts defined as ≥10 consecutive purines ([AG]{10,}). Candidate tracts were ranked by proximity to the LTR boundary, and the closest match was reported as the PPT.

#### Evidence aggregation and confidence scoring

Each LTR was assigned an overall confidence score computed as the sum of the promoter score, PAS score, and binary contributions for detected PBS and PPT signals. A consolidated evidence string reports detected promoter motifs, final R and U5 lengths, PAS features (PAS hexamer, CA cleavage site, downstream GU-rich element), and PBS/PPT matches.
