## Supplemental Results for "Regulatory Features and Functional Specialization of Human Endogenous Retroviral LTRs: A Genome-Wide Annotation and Analysis via HERVarium"

### Supplementary Results — Benchmarking the U3–R–U5 Segmentation Using Shuffled LTR Controls

To evaluate whether the reconstructed U3–R–U5 structures reflect non-random retroviral organisation rather than chance motif co-occurrence, we compared the confidence scores of real LTRs to those obtained from a shuffled negative control. For each LTR, a length- and base-composition–matched randomized sequence was generated by permuting nucleotide order, thereby preserving overall compositional bias while disrupting promoter and cleavage-site grammar.

Both real and shuffled sequences were processed using the full U3–R–U5 segmentation workflow. Because the confidence score integrates promoter evidence, PAS support, and geometry constraints, score distributions provide a non-thresholded measure of structural coherence.

Real LTRs exhibited higher confidence scores than shuffled controls, accompanied by a clear shift in the score distribution (Wilcoxon rank-sum test with continuity correction, $p < 2.2 \times 10^{-16}$).

Together, these analyses indicate that the U3–R–U5 architectures recovered by the segmentation pipeline reflect non-random retroviral organisation and are unlikely to arise from motif frequency alone.
